# Sex disparities in age-related neuromuscular decline: unveiling female susceptibility from early to late elderly

**DOI:** 10.1101/2023.06.13.544761

**Authors:** Yuxiao Guo, Eleanor J. Jones, Thomas F. Smart, Abdulmajeed Altheyab, Nishadi Gamage, Daniel W. Stashuk, Jessica Piasecki, Bethan E. Phillips, Philip J. Atherton, Mathew Piasecki

## Abstract

**Background:** Females typically have a longer lifespan than males which is not matched by an improved healthspan, with older females having higher rates of frailty, characteristic of a sex specific degradation of the neuromuscular system. Several motor unit (MU) characteristics show sex-specific behaviour during mid-level contractions in healthy younger people, highlighting a potential influence of hormonal differences that may be augmented in older age. The purpose of this study was to investigate sex differences in physical performance and MU features of the aged human vastus lateralis (VL) from early to late elderly.

**Methods:** This study included 21 healthy older males (mean ± SD, range: 67.2 ± 7.6, 56 – 81 yrs) and 17 healthy older females (69.5 ± 5.2, 60 – 78 yrs). Intramuscular electromyography data were collected from VL during standardised submaximal sustained contractions. Muscle size and physical performance characteristics were also measured. Multiple mixed-effects linear regression models with age considered were conducted and statistical significance was accepted when p<0.05.

**Results:** When compared to males, early to late elderly females had smaller cross-sectional area of VL (p<0.001), lower knee extensor torque (p<0.001) and poorer force steadiness (p=0.036), as well as higher MU firing rate (FR) (p=0.025) and greater MU FR variability (p=0.031). With progression from early to late elderly, both sexes showed decreased functional capacity at a similar rate.

**Conclusion:** Functional deterioration occurs to a similar extent in both sexes from early to late elderly. However, throughout the majority of the elderly period males demonstrate a greater muscle size, strength, and functional performance. Older females have greater MUFR variability and worse force steadiness than older males. These findings help to address the lack of MU data in older females, and suggest earlier interventions are needed in older females to prevent functional deterioration and reduce the health-sex paradox within ageing humans.

## Introduction

Sex differences in the ageing process are well established with females typically living longer, yet do so in poorer health ^1^. Though ageing is generally characterized by decreased muscle mass and strength, there is a significant heterogeneity with respects to physical function. Despite living longer, females are reported to have lower levels of physical performance with a higher frailty index score throughout the lifespan when compared to males, which is exacerbated with increasing age ^2^. Yet there is limited mechanistic data on neuromuscular function in older females. Thus, there is a growing need to better understand the underlying reasons for these sex differences in aging and their implications for health outcomes to generate better informed interventions.

Among the most underexplored elements accounting for sex-specific physical performance are alterations of motor unit (MU) properties. Voluntary contraction of muscle relies upon the coordinated activation of MUs, sets of muscle fibres innervated by a single motoneuron ^3^, which can be recruited based on their size and further modulated via the frequency at which they discharge, referred to as MU recruitment and firing rate (FR) modulation, respectively ^4^. There were ∼30% fewer MUs in the quadriceps and tibialis anterior muscles of older males compared to younger males ^5,6^, which contributes to loss of functionality in older age. Following denervation, a muscle fibre may atrophy and eventually be lost, or it may be ‘rescued’ by an adjacent surviving axon and the formation of a new neuromuscular junction (NMJ). This process of MU remodelling results in aged MUs that are fewer in number but larger in size ^7^. These MU adaptations may alter neuromuscular recruitment strategies and/or further influence force control ^8^. However, most of the knowledge about neuromuscular function in VL are primarily available in males and/or a smaller number of young females, with a distinct lack of direct comparisons.

Sex steroids are key players in the aging process and have been identified as potential contributors to biological sex differences in neuromuscular function ^9^. The most common groups of sex steroids include androgens (such as testosterone), which are typically more abundant in males. Testosterone has an anabolic impact on skeletal muscle and its precursor dehydroepiandrosterone (DHEA) was positively associated with MUFR in highly active and inactive older males ^10^. Estrogens and progestogens are the predominant female sex hormones and have the ability to cross from the blood brain barrier potentially influencing crucial central nervous system function ^11^. The decline in estrogen and androgen levels with aging has been shown to contribute to the enhanced decline in muscle function, as both exhibit a range of neuroprotective effects in motoneurons, such as dendritic maintenance and axonal sprouting ^12^. Additionally, the dramatic decline in estrogen levels following the menopause heightens the risk of osteoporosis for many older females ^13^, further impacting on physical function and mobility, and increasing the risk of fracture compared to males.

We have previously reported a significant decline in MU FR in tibialis anterior from middle to older highly active females, which was not observed in highly active males ^14^, and separately, a higher MU FR in vastus lateralis (VL) in healthy young females comparatively to males at normalised force levels ^15^. This highlights a potential influence of hormonal differences, which may be amplified in older age. The quadriceps muscles are highly susceptible to atrophy in aging and are closely related to functional decline ^16^, therefore, the purpose of the present study was to explore the influences of sex and ageing on physical performance and neuromuscular characteristics in VL of early to late elderly males and females. It was hypothesized that females would have lower functional performance than males, and markers of more extensive MU remodelling than males.

## Methods

### Participants

This study was approved by the University of Nottingham Faculty of Medicine and Health Sciences Research Ethics Committee (90-0820, 407-1910, 390-1121) and was conducted between 2019-2022 in accordance with the Declaration of Helsinki.

Twenty-one healthy males and seventeen healthy females between the ages of 55 to 85 years were recruited from the local community via advertisement. All recruited participants were recreationally active on a day-to-day basis. Prior to enrolment, all participants completed a comprehensive clinical screening examination based at the School of Medicine, Royal Derby Hospital Centre and subsequently provided written informed consent. Screening procedure allowed exclusion of the participants with musculoskeletal abnormalities, acute cerebrovascular or cardiovascular diseases, uncontrolled hypertension, or metabolic disease.

### Anthropometry

Prior to testing, calibrated scales and a stadiometer were used to assess the body mass and height of each participant for calculation of the body mass index. Cross-sectional area (CSA) of the VL was measured using an ultrasound probe (LA532 probe B-mode, frequency range 26-35 Hz and MyLab™50 scanner, Esaote, Genoa, Italy). To quantify VL CSA, images were captured from the aponeurosis borders of the VL of the right leg in a medial-lateral fashion using panoramic imaging (VPAN) and further analysed with ImageJ software (National Institute of Health, USA) by tracing around the VL following the contour of the aponeurosis. CSA was determined by analysing each image three times and taking the average of three images.

### Balance

A Footscan plate (Footscan, 200 Hz, RScan International, Belgium) was used to assess the postural sway during right-leg standing with eyes open. Participants were asked to step on the platform and visually focus on a target point in front of them for the duration of the test (10 seconds). A 5-second countdown was given before instruction to lift the left leg. Travelled distance and ellipse area of centre of pressure (COP) were recorded. Each participant was allowed three attempts and the best value was reported. The total distance travelled by COP during the test time was determined as COP travelled distance in millimetres (mm). COP ellipse area was calculated based on the data in the ellipse set for 95% of the total area covered by the COP trajectory in both anterior-posterior and medial-lateral directions. Smaller area and/or travelled distance indicated a better postural balance.

### Grip Strength

A handgrip dynamometer (Grip-D, Takei, Japan) was used to test the grip strength of the dominant hand. When measuring grip strength, the dynamometer was held in a standing position with the pointer facing outward, and the width of the grip was adjusted so that the index finger’s interphalangeal joint was bent 90 degrees. The arms were lowered naturally, the feet were hip width apart, and the dynamometer was grasped with maximum force without touching the body or clothing. The maximal value among three measurements was accepted as the grip strength.

### Timed Up and Go (TUG)

The TUG test was used to evaluate dynamic balance and functional mobility. TUG test involved getting up from a standard chair without armrests, walking for 3 metres, turning around an obstacle and returning to the original sitting position as quickly as possible without running. The test was initiated with a verbal “go” instruction from the researcher, and the time taken to complete the test was recorded. After a familiarisation attempt, the participant was asked to have a real go after one minute rest.

### Knee Extensor Torque

Participants were placed in a custom-built chair which was altered to ensure the hips and knees were flexed at ∼90□. The right lower leg was attached to a force dynamometer with a non-compliant strap (purpose-built calibrated strain gauge, RS125 Components Ltd, Corby, UK) above the medial malleolus. The hips and pelvis were also securely held by a seat belt to reduce movement of the hips and upper trunk during contractions. The distance between the centre of the force strap and the lateral femoral condyle was measured to determine the external knee joint moment arm. With real-time visual feedback and verbal encouragement, participants were instructed to perform each trial with maximal effort as hard and fast as they could following a standardized warm-up of submaximal contractions. During the trial, participants were not allowed to hold onto the side of the chair and were asked to cross their arms across the chest. It was further repeated two to three times with 60 seconds rest intervals between each one. If there was a difference of less than 5% between the two last attempts, the highest value was accepted as the maximal voluntary contraction (MVC). Torque was then determined by multiplying the selected MVC by the lever arm. To quantify force steadiness during 10% and 25% of MVC, the coefficient of variation of the force (CoV) was calculated = (SD/Mean)*100. When calculating CoV, in an attempt to reduce corrective actions, the first two passes of the target (<1s) were excluded from the analysis.

### Intramuscular Electromyography (iEMG)

Prior to intramuscular needle electrode insertion, a familiarisation trial was performed in which the participant had an attempt to contract to each submaximal target line at 10% and 25% of MVC observed on a monitor. Following this, a 25-mm or 40mm disposable concentric needle electrode (N53153; Teca, Hawthorne, New York, USA) was inserted into the muscle belly of VL of the right leg, to a depth of 1.5-2 cm depending on muscle size. A ground electrode was placed over the patella of the same testing leg. To ensure an adequate signal to noise ratio once the needle was positioned, the participants were asked to perform several low-level voluntary contractions. Participants were then asked to perform sustained voluntary isometric contractions at 10 % and 25 % of MVC, 4 times each, with 20-second intervals between contractions. Each time, the needle was repositioned 180 degrees by twisting the bevel edge and withdrawing 5 mm each time to sample from spatially distinct areas ^17^. iEMG signals were recorded in Spike2 (Version 9.06), sampled at 50 kHz and bandpass filtered at 10 Hz to 10 kHz (1902 amplifier; Cambridge Electronic Design Ltd, Cambridge, UK) and stored for future off-line analysis.

### Intramuscular Electromyography Analysis

Using decomposition-based quantitative electromyography (DQEMG) software, individual motor unit potentials (MUPs) were detected, and motor unit potential trains (MUPTs) were extracted from the iEMG signal recorded during the sustained force portion of the contraction, enabling the evaluation of VL electrophysiological activity during contractions. MUPs were visually inspected and markers corresponding to MUP onset and end were adjusted if required. MU FR is reported as the estimated number of MU discharges per second (Hz) within a MUPT. MU FR variability is reported as the estimate of coefficient of variation (CoV) of the inter-discharge intervals (IDIs), displayed as a percentage. MUP area was determined as the total area within the MUP duration (onset to end). A MUP’s complexity is determined by the number of phases, which are defined as the number of components above or below the baseline.

An near fibre MUP (NFM) is calculated by applying a low-pass second-order differentiator to a MUP. In this way, contributions from fibre near to the needle electrode primarily contribute to a NFM and interference from distant active fibres is reduced. All NFMs (and corresponding MUPs) without distinct peaks were excluded from analyses. NFM jiggle is a measure of the shape variability of consecutive NFMs of an MUPT expressed as a percentage of the total NFM area ^18^.

### Statistics

All statistical analysis was conducted using R Studio (Version 2022.07.1). Descriptive data were generated for all variables. *Multiple linear regression models* with age considered as a covariate were used to compare the effects of age and sex on physical characteristics. *Multilevel linear regression mode*ls using the *lme4* package (Version 1.1-27.1) ^19^ with age considered as a co-variate were generated to compare MU parameters between groups across two contraction levels in order to preserve variability between and within participants simultaneously. Sampled MUs were regarded as the first level of multilevel models, and participants with their respective clustered sets of sampled MUs were regarded as the second level. The results are displayed as coefficient estimates, 95% confidence intervals and p values. Standardised estimates were calculated through the package *effectsize* (Version 0.8.2) ^20^ for forest plotting. Statistical significance was accepted when *p*<0.05.

## Results

Thirty-eight older participants were included in the study, consisting of 21 healthy older males (age range: 56-81 years) and 17 healthy older females (age range: 60-78 years). All recorded measures are shown in Table 1.

**Table 1.**
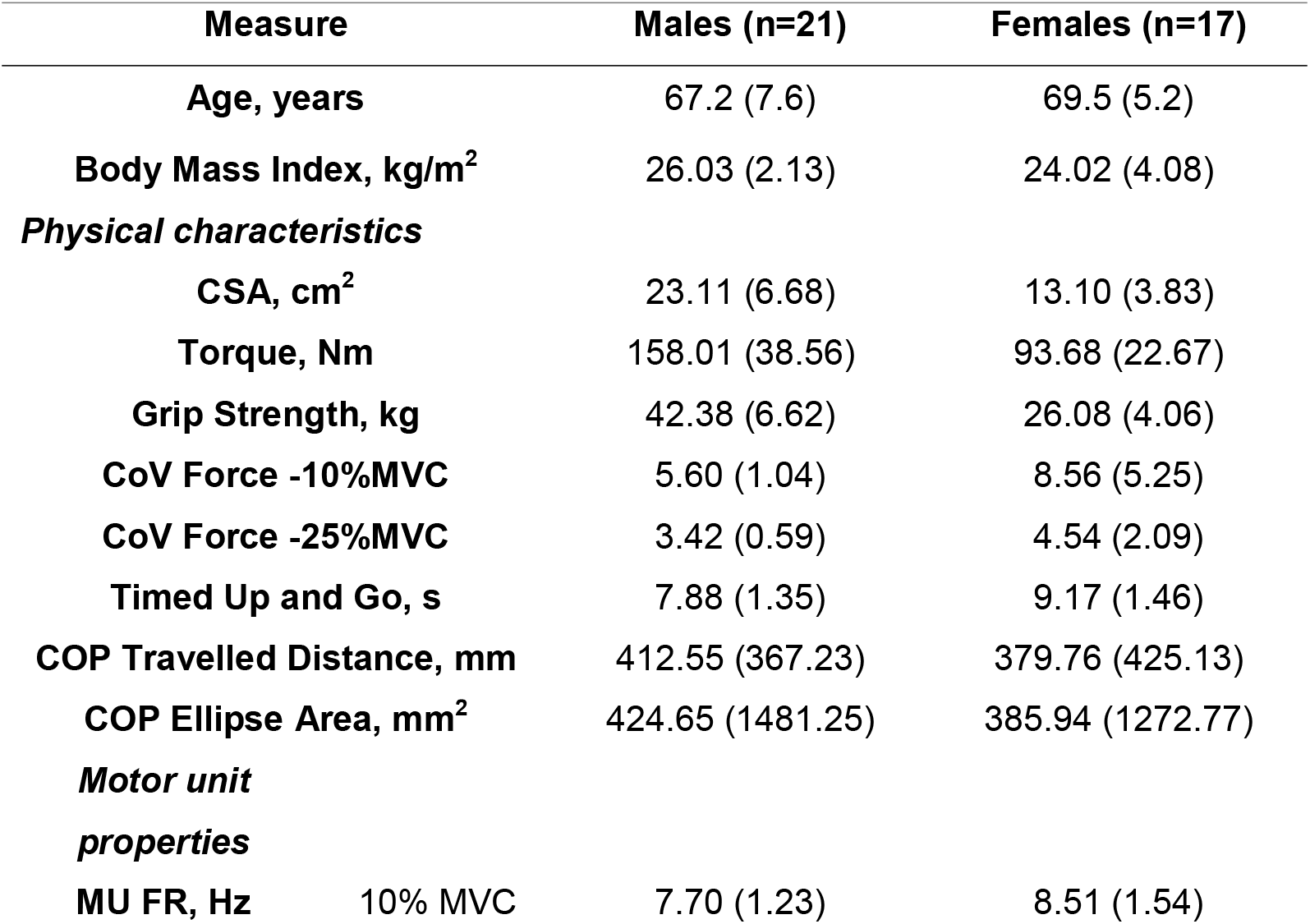
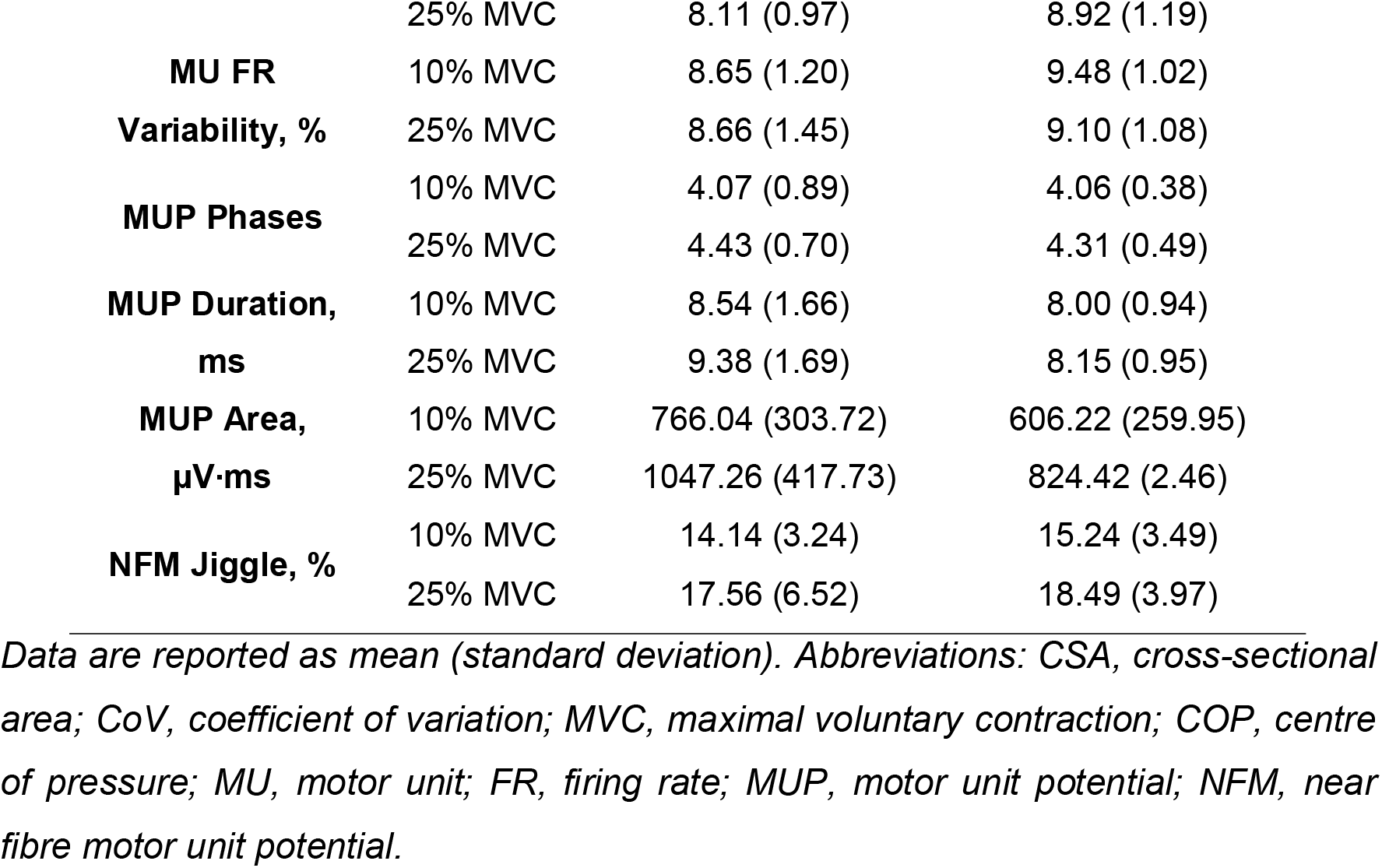
Participant characteristics.

There were no significant interactions between sex and age detected in functional parameters. When adjusting for sex, for every year increase in age, muscle cross-sectional area decreased by 0.45 cm^2^ (95%CI: -0.70 to -0.21; p=0.001) and muscle torque decreased by 1.62 Nm (−3.20 to -0.04; p=0.045). There was no significant difference in grip strength with advancing age (−0.27; -0.58 to 0.04; p=0.084). Though non statistically significant, for every year increase in age, TUG increased by 0.07 s (−0.002 to 0.14; p=0.058). Additionally, force steadiness decreased significantly at both 10% (beta: 0.09; 0.03 to 0.15; p=0.003) and 25% MVC. (0.04; 0.002 to 0.07; p=0.041) and unilateral COP ellipse area increased (7.93; 1.82 to 14.04; p=0.013). However, there was no significant difference in unilateral COP travelled distance (6.54; -2.33 to 15.40; p=0.143).

When adjusting for age, females had 55.3% smaller muscle cross-sectional area (- 8.93; -12.23 to -5.62; p<0.001), 48.1% lower knee extensor torque (−60.58; -81.49 to -39.68; p<0.001), 44.7% lower grip strength (−15.60; -19.75 to -11.44; p<0.001), and 15.1% longer TUG times (1.14; 0.23 to 2.04; p=0.015) than males. Females showed 41.8 and 28.1% poorer force steadiness at 10% (1.58; 0.79 to 2.37; p<0.001) and 25% MVC (0.59; 0.13 to 1.06; p=0.014) when compared to males. However, there were no sex differences in unilateral COP travelled distance (−48.05; -161.48 to 65.38; p=0.394) or COP ellipse area (−44.02; -122.67 to 34.63; p=0.263). All statistical model outputs are shown in Table 2 and individual data are shown in Figure 1.

**Table 2.** Summary of multiple linear regression analysis for functional parameters.

|  | Age <sup>a</sup> |  |  | Sex <sup>b</sup> |  |  |
| --- | --- | --- | --- | --- | --- | --- |
|  | beta | 95% CI | p value | beta | 95% CI | p value |
| CSA, cm<sup>2</sup> | -0.45 | -0.70 to -0.21 | **0.001** | -8.93 | -12.23 to -5.62 | **<0.001** |
| Torque, Nm | -1.62 | -3.20 to -0.04 | **0.045** | -60.58 | -81.49 to -39.68 | **<0.001** |
| Grip Strength, kg | -0.27 | -0.58 to 0.04 | 0.084 | -15.60 | -19.75 to -11.44 | **<0.001** |
| Timed Up and Go, s | 0.07 | -0.002 to 0.14 | 0.058 | 1.14 | 0.23 to 2.04 | **0.015** |
| CoV Force- 10% MVC | 0.09 | 0.03 to 0.15 | **0.003** | 1.58 | 0.79 to 2.70 | **<0.001** |
| CoV Force – 25% MVC | 0.04 | 0.002 to 0.07 | **0.041** | 0.59 | 0.13 to 1.06 | **0.014** |
| COP Travelled Distance, mm | 6.54 | -2.33 to 15.40 | 0.143 | -48.05 | -161.48 to 65.38 | 0.394 |
| COP Ellipse Area, mm<sup>2</sup> | 7.93 | 1.82 to 14.04 | **0.013** | -44.02 | -122.67 to 34.63 | 0.263 |

<sup>a</sup> Multiple linear regression model with **Age** as dependent variable; <sup>b</sup> multiple linear regression model with **Sex** as dependent variable. Values in bold reflect statistically significant ( $p < 0.05$ ) results. Abbreviations: CI, confidence interval; CSA, cross-sectional area; CoV, coefficient of variation; MVC, maximal voluntary contraction; COP, centre of pressure.

**Table 3.** Summary of linear regression analysis for motor unit parameters.

|  | Sex <sup>a</sup> |  |  | Level <sup>b</sup> |  |  |
| --- | --- | --- | --- | --- | --- | --- |
|  | beta | 95% CI | p value | beta | 95% CI | p value |
| MU FR, Hz | 0.91 | 0.15 to 1.68 | **0.025** | 0.37 | 0.17 to 0.58 | **<0.001** |
| MU FR<br>Variability, % | 0.89 | 0.10 to 1.68 | **0.031** | 0.33 | -0.11 to 0.77 | 0.144 |
| MUP Phases | -0.07 | -0.48 to 0.33 | 0.731 | 0.26 | 0.10 to 0.42 | **0.001** |
| MUP Duration, ms | -0.76 | -1.60 to 0.08 | 0.084 | 0.64 | 0.34 to 0.93 | **<0.001** |
| MUP Area, $\mu\text{V}\cdot\text{ms}$ | -217.59 | -420.40 to -<br>14.78 | **0.042** | 246.68 | 181.63 to<br>311.73 | **<0.001** |
| NF Jiggle, % | 0.10 | -3.05 to 3.26 | 0.949 | 2.21 | 0.71 to 3.72 | **0.004** |

<sup>a</sup> Multiple mixed-effects linear regression model with sex as dependent variable; <sup>b</sup> multiple mixed-effects linear regression model with contraction level as dependent variable. Beta value and 95% confidence interval (CI) represent the model predicted change per unit between sexes and contraction levels with age considered as a covariate. Values in bold reflect statistically significant ( $p < 0.05$ ) results. Abbreviations: MU, motor unit; FR, firing rate; MUP, motor unit potential; NFM, near fibre motor unit potential.

**Figure 1.**
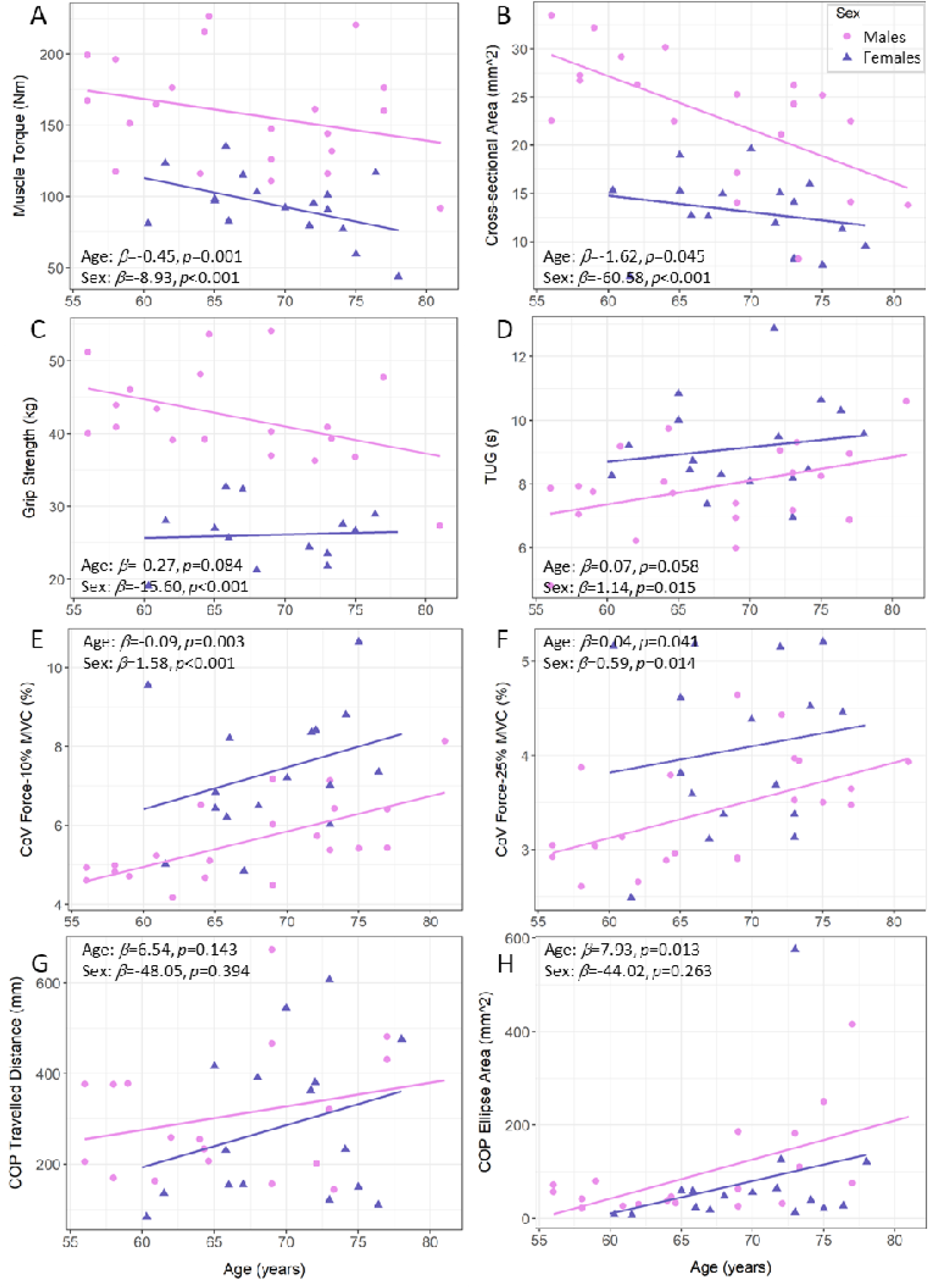
Scatter points of physical parameters in older males (pink circles) and females (purple triangles). The estimates of the main effect of Age and Sex from multiple linear regression models for physical parameters with no interaction detected are reported in text boxes. For data visualisation, male and female regression lines are displayed separately. Abbreviations: TUG, timed up and go; CoV, coefficient of variation; MVC, maximal voluntary contraction; COP, centre of pressure.

**Figure 2.**
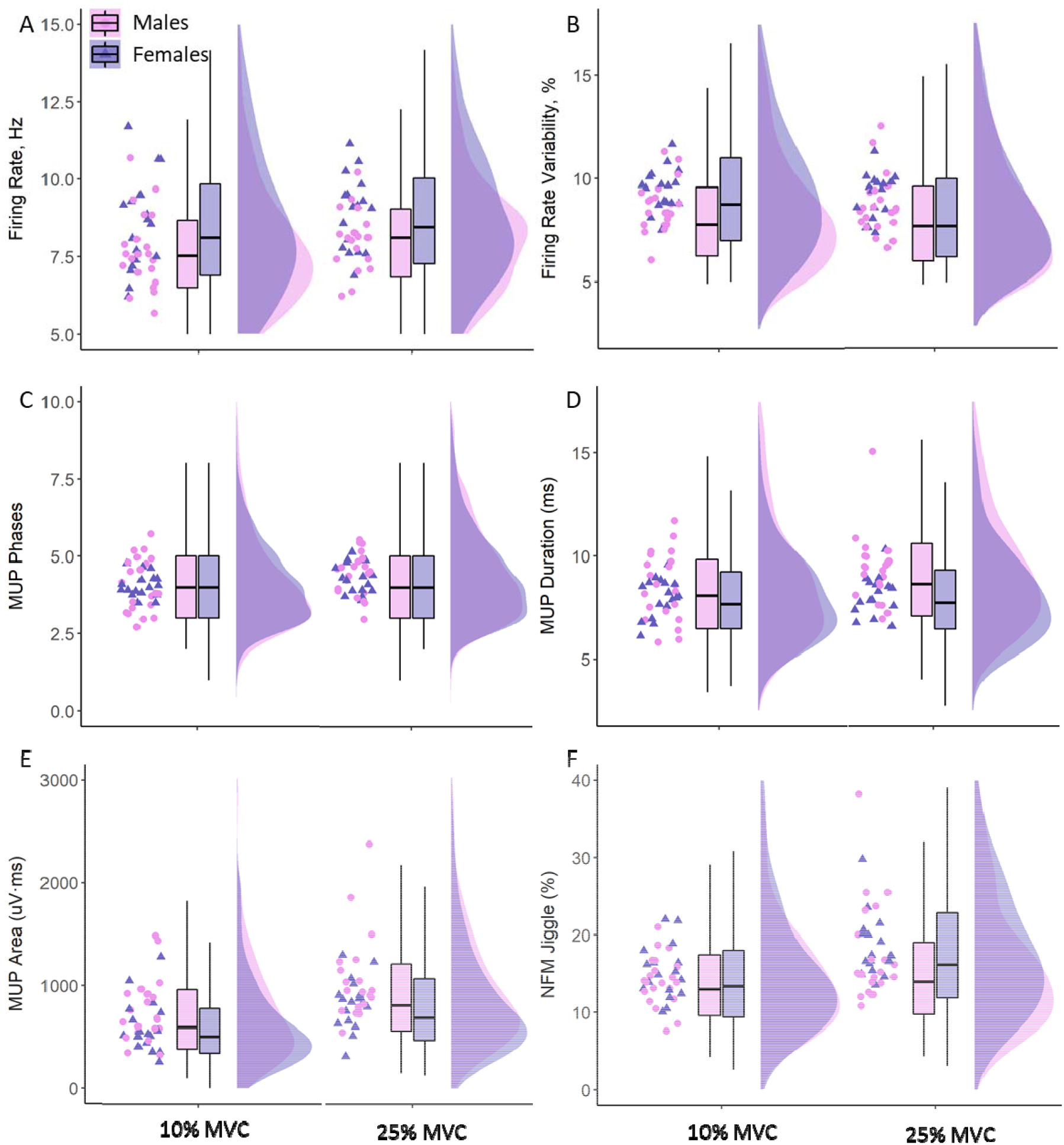
Motor unit (MU) properties in males (pink) and females (purple) at 10% and 25% maximal voluntary contraction (MVC). Individual participant means are shown in the left column within each plot with males in circles and females in triangles. Box plots illustrate the first and third quartiles, and the median of all MUs. Distributions of all MUs are also shown in each density plot.

**Figure 3.**
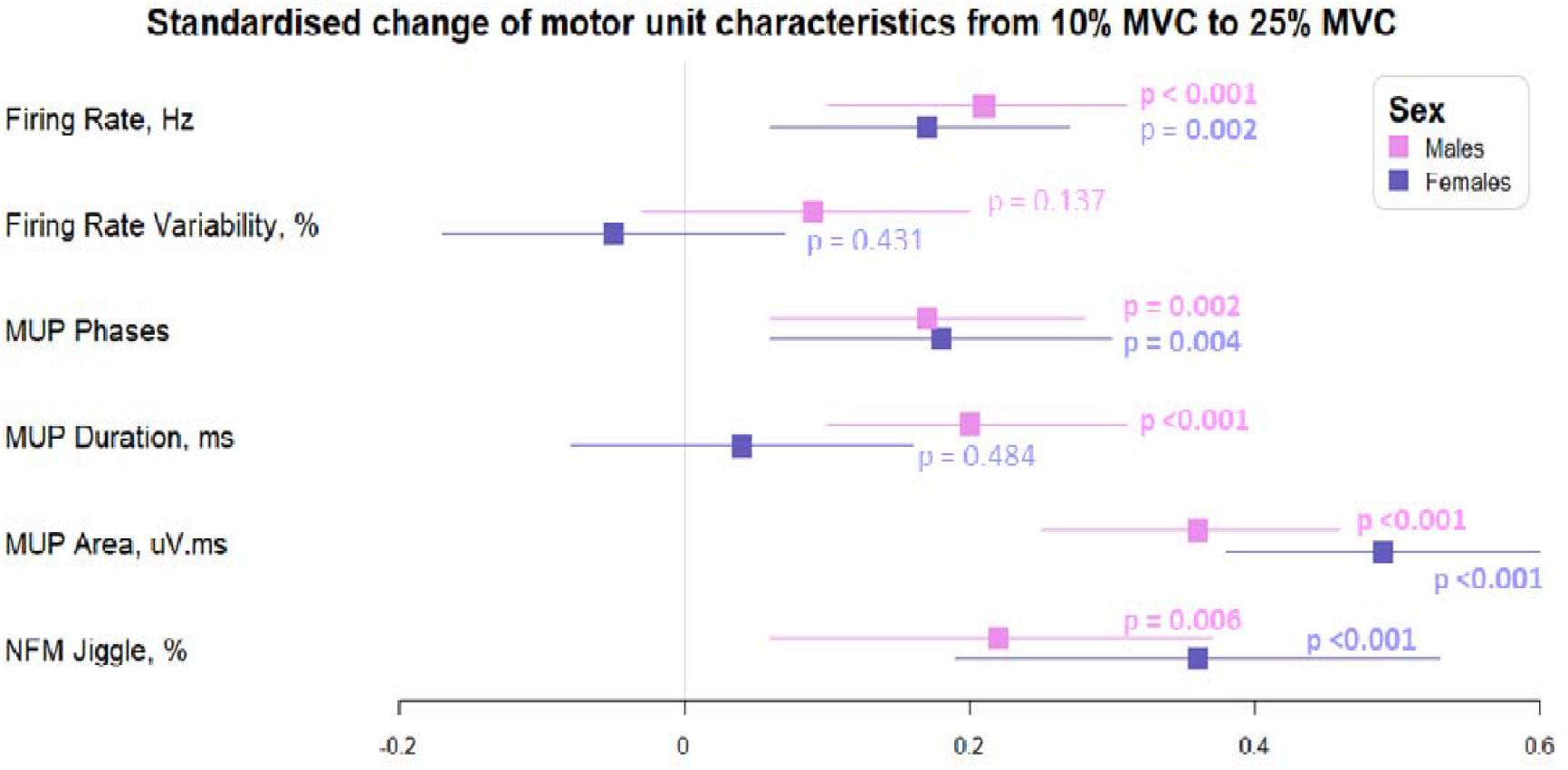
Forest plot of the standardised regression coefficient estimate for associations between motor unit characteristics and contraction levels in older males (pink) and females (purple). Beta and 95% confidence intervals represent the standardised model predicted change per unit from 10% to 25% maximal voluntary contraction (MVC). All statistical analysis was based on multilevel mixed effects linear regression model with age considered as a covariate. Standardised values for each parameter make the comparisons between older males and females justifiable.

At 10% MVC, the mean number of sampled Mus per person was 21 in males with a mean of 5 Mus sampled per needle position, and 24 in females with 6 Mus sampled per needle position. At 25% MVC, the mean number of Mus sampled per person was 33 in males with 8 Mus sampled per needle position and 38 in females with 9 Mus sampled per needle position.

There was no association between age and all MU parameters after adjusting for sex and contraction level (all p>0.05). There was no significant sex x contraction level interaction for FR (p=0.955) but there was a main effect of sex (p=0.025) with females showing a greater FR when compared to males. MUFR increased significantly when moving from low- to mid-level contractions (p<0.001) in both sexes. Similarly, there was no significant sex x contraction level interaction for FR variability (p=0.115) but there was a main effect of sex (p=0.031) with females showing a higher FR variability when compared to males. There was no significant difference in FR variability when moving from low- to mid-level contraction (p=0.144).

There was no significant sex x contraction level interaction for MUP phases (p=0.782), and there was no statistical difference between sexes (p=0.731) whereas there was a main effect of contraction level, with a greater number of MUP phases observed at 25% when compared to 10% MVC contractions (p=0.001). A significant sex x contraction level interaction was observed in MUP duration (p=0.014) with females exhibiting a shorter MUP duration at mid-level contractions (p<0.001). There was no significant sex x contraction interaction for MUP area (p=0.973) but there was a main effect of sex (p=0.042), with a smaller MUP area in females when compared to males, and both sexes showing greater MUP area when moving from low to mid-level contractions (p<0.001).

There was no significant sex and contraction level interaction for NF jiggle (p=0.371) and no significant main effect of sex (p=0.949) however, NF jiggle increased significantly when moving from low to mid-level contractions (p=0.004).

## Discussion

The present study demonstrated an age-related decrease in physical performance in early to late elderly humans, occurring at a similar rate in males and females, yet with distinct sex differences. In older males and females, muscle size, muscle strength, force steadiness and unilateral balance ability decreased with increasing age. Regardless of the progressive decrease in both sexes, females had a smaller muscle size, lower knee extensor and grip strength, longer TUG time and poorer force steadiness than males. iEMG derived MU data from the VL show females exhibited smaller markers of MU size, higher MU FR and greater MU FR variability. When assessing changes in the characteristics of active MUs from low- to mid-level contractions, both males and females showed higher MU FR, MUP phases, MUP area and NF jiggle. These data underscore the connection between diminished functionality in older females and a lower baseline observed during early stages of aging, potentially serving as a valuable reference for pinpointing pivotal inflection points along the path towards frailty.

Numerous studies have highlighted the age-related reduction of muscle size and strength ^21,22^ and the quadriceps appear to be particularly susceptible when compared to other lower extremity muscles such as hamstrings and tibialis anterior ^23^. Interestingly, the ∼55% lower muscle size and ∼48% lower strength in older females reported here is greater than the sex-based differences we have previously reported when comparing healthy young (∼39% and ∼37%, respectively) ^15^, and suggests the sex-based disparities are augmented in older age, to the detriment of females.

Grip strength has been identified as a robust predictor of frailty and mortality among older individuals. Our data are in alignment with large-scale investigations ^24^ that identified lower grip strength in females when compared to males in early and late older age. However, our multiple regression model did not exhibit a significant decline in grip strength from early to late elderly. This discrepancy may be attributed to the smaller sample size in this cohort. Furthermore, it has been suggested that females exhibit better preservation of grip strength than males throughout the aging process ^25^, implying that diminished grip strength alone cannot fully predict or elucidate systemic physical frailty ^26^. Similarly, even though age did not seem to have a statistically significant effect, sex-based differences in TUG scores were observed with females showing higher scores than males, indicating that females in their early and late older age have poorer physical performance.

Force fluctuations at both low- and mid-level submaximal sustained contractions in both sexes are greater in old compared to young ^27,28^ and here we show a continued decrease in force steadiness from early to late elderly. This age-related loss is likely multifactorial and includes altered synaptic input to MUs, altered MU discharge properties ^29,30^, as well as the loss and subsequent remodelling of MUs ^7,31^. Additionally, considering multiple muscles are involved in the knee extension movements, an increased co-activation of agonist and antagonist muscles in lower limb has been reported with age and may play an important role in regulating force and control ability ^32,33^.

The hormonal milieu is one of the most distinguishing features of the two sexes, with the predominant female sex hormone estrogen and progesterone fluctuating across a normal menstrual cycle, whilst the male sex hormone testosterone remains relatively constant. In crossing the blood brain barrier, estrogen elicits excitatory effects via the potentiation of glutamatergic receptors ^34^ while progesterone increases activity of gamma-aminobutyric acid (GABA), causing inhibitory effects ^11,35^. Following the menopause and the cessation of the menstrual cycle, typically occurring at the age of ∼51 yrs ^36^, estrogen concentrations plummet and generate a severe change in the hormonal milieu that is not apparent in males. The exclusive manifestation of this pronounced change in females may possibly generate an impaired equilibrium between neuronal excitation and inhibition in advanced age. Alterations in synaptic plasticity have been linked to reduced estrogen levels, albeit in animal models ^37,38^, and a significantly lower level of GABA signalling components has also been reported in the cortex of older females ^39^. This disruption may culminate in increased MUFR variability and an exacerbation of the decline in force steadiness, as observed here in older females during low- and mid-level contractions. Notably, this sex difference was not apparent from our prior findings from the same muscle in younger cohorts ^15^.

In addition to greater MU FR variability, females also had higher MU FRs than males, accompanied by smaller MUP areas. This is consistent with what we have previously observed in the same muscle in younger cohorts ^15^, and although not definitive, it may indicate that at a given force level females recruit smaller MUs which discharge at higher rates, and this strategy is maintained in older age. With force increases, we found that recruitment strategies for early and late elderly males and females did not differ, also reported for young cohorts ^15^. Put simply, when moving from a low to a mid-level contraction both sexes employed both MU rate coding and recruitment, as indicated by higher MU FR and larger MUP area, to a similar extent.

The duration of a MUP is largely dependent on the number of fibres contributing the potential and the temporal dispersion of their depolarisation ^18^. Here, there was a sex and level interaction for MUP duration indicating distinct adaptations between males and females when moving from low-to mid-level contractions. Males exhibited extended MUP duration whereas no distinct difference was observed in females. This disparity may be attributed to the recruitment of MUs with greater numbers of muscle fibres in males, or a greater increase in the temporal dispersion of their depolarisation as a result of a larger MU territory. To better understand nerve-muscle communication, we explored near fibre jiggle which is indicative of NMJ transmission instability ^18^. There was no significant sex-difference in NF Jiggle, but an increased NF jiggle was detected in both males and females when moving from low-to mid-level contractions.

## Strengths and Limitations

This is the first study using iEMG techniques to explore VL MU characteristics of early to late older males and females, enabling the avoidance of signal attenuation due to greater subcutaneous tissues in older females. This direct sex comparison contributes to the lack of data on older females and reveals some of the functional detriments are also accompanied by greater MU adaptation. Indeed, the sex difference of muscle size and strength exceeds that which we have previously reported in healthy young. However, we have sampled MU activities at low- and mid-level contractions and these findings cannot be extrapolated to Mus recruited at higher contraction intensities. Secondly, the right leg was uniformly assessed across all participants and as such, cannot infer possible bilateral differences which may contribute to functional impairment.

## Conclusions

Functional deterioration progresses at a similar rate in both sexes from early to late elderly. Males demonstrate a greater muscle size and strength, better motor control and functional performance. Females have greater MUFR and MUFR variability and poorer force steadiness than males, a sex-specific trait that was not apparent in younger cohorts. Moreover, the greater sex-based differences in aged muscle may contribute to greater functional declines in older females. These findings add to the paucity of data in older females and suggest early interventions are needed for older females to prevent functional deterioration.

## Acknowledgements

We thank all of the participants for their enthusiastic involvement in this study.

## Author contributions

YG, JP, PJA, BEP and MP contributed to the conception and design of the work. YG, EJJ, TFS AA and NG acquired the data. YG, EJJ, TFS, and AA analysed the data. YG and MP drafted the manuscript and prepared the figures. YG and MP contributed to the interpretation of the results. All authors contributed to the revision of the manuscript. All authors have approved the final version of the submitted manuscript for publication and are accountable for all aspects of the work. All persons designated as authors qualify for authorship, and all those who qualify for authorship are listed.

## Data Availability Statement

The datasets generated and analysed during the current study are available from the corresponding author upon reasonable request.

## References

1. Carmel S. Health and Well-Being in Late Life: Gender Differences Worldwide. Front. Med.. 2019;6.

2. Gordon EH, Peel NM, Samanta M, Theou O, Howlett SE, Hubbard RE. Sex differences in frailty: A systematic review and meta-analysis. Exp Gerontol 2017;89:30–40.

3. Heckman CJ, Enoka RM. Motor unit. Compr Physiol 2012;2:2629–2682.

4. Enoka RM, Duchateau J. Rate Coding and the Control of Muscle Force. Cold Spring Harb Perspect Med 2017;7.

5. Piasecki M, Ireland A, Stashuk D, Hamilton-Wright A, Jones DA, McPhee JS. Age-related neuromuscular changes affecting human vastus lateralis. J Physiol 2016;594:4525–4536.

6. Piasecki M, Ireland A, Piasecki J, Stashuk DW, Swiecicka A, Rutter MK et al. Failure to expand the motor unit size to compensate for declining motor unit numbers distinguishes sarcopenic from non-sarcopenic older men. J Physiol 2018;596:1627–1637.

7. Jones EJ, Chiou S-Y, Atherton PJ, Phillips BE, Piasecki M. Ageing and exercise-induced motor unit remodelling. J Physiol 2022;600:1839–1849.

8. Pethick J, Piasecki M. Alterations in Muscle Force Control With Aging: Is There a Modulatory Effect of Lifelong Physical Activity?. Front. Sport. Act. Living. 2022;4.

9. Hunter SK. Sex differences in human fatigability: mechanisms and insight to physiological responses. Acta Physiol (Oxf) 2014;210:768–789.

10. Guo Y, Piasecki J, Swiecicka A, Ireland A, Phillips BE, Atherton PJ et al. Circulating testosterone and dehydroepiandrosterone are associated with individual motor unit features in untrained and highly active older men. GeroScience 2022;44:1215–1228.

11. Del Río JP, Alliende MI, Molina N, Serrano FG, Molina S, Vigil P. Steroid Hormones and Their Action in Women’s Brains: The Importance of Hormonal Balance. Front. Public Heal.. 2018;6.

12. Hyer MM, Phillips LL, Neigh GN. Sex Differences in Synaptic Plasticity: Hormones and Beyond. Front. Mol. Neurosci.. 2018;11.

13. Sözen T, Özışık L, Başaran NÇ. An overview and management of osteoporosis. Eur J Rheumatol 2017;4:46–56.

14. Piasecki J, Inns TB, Bass JJ, Scott R, Stashuk DW, Phillips BE et al. Influence of sex on the age-related adaptations of neuromuscular function and motor unit properties in elite masters athletes. J Physiol 2021;599:193–205.

15. Guo Y, Jones EJ, Inns TB, Ely IA, Stashuk DW, Wilkinson DJ et al. Neuromuscular recruitment strategies of the vastus lateralis according to sex. Acta Physiol 2022;235:e13803.

16. Naruse M, Trappe S, Trappe TA. Human skeletal muscle-specific atrophy with aging: a comprehensive review. J Appl Physiol 2023;134:900–914.

17. Jones EJ, Piasecki J, Ireland A, Stashuk DW, Atherton PJ, Phillips BE et al. Lifelong exercise is associated with more homogeneous motor unit potential features across deep and superficial areas of vastus lateralis. GeroScience 2021;43:1555–1565.

18. Piasecki M, Garnés-Camarena O, Stashuk DW. Near-fiber electromyography. Clin Neurophysiol Off J Int Fed Clin Neurophysiol 2021;132:1089–1104.

19. Bates D, Mächler M, Bolker B, Walker S. Fitting Linear Mixed-Effects Models Using lme4. J Stat Softw 2015;67:1–48.

20. Ben-Shachar M, Lüdecke D, Makowski D. effectsize: Estimation of Effect Size Indices and Standardized Parameters. J Open Source Softw 2020;5:2815.

21. Piasecki M, Ireland A, Stashuk D, Hamilton-Wright A, Jones DA, McPhee JS. Age-related neuromuscular changes affecting human vastus lateralis. J Physiol 2016;594:4525–4536.

22. Wilkinson DJ, Piasecki M, Atherton PJ. The age-related loss of skeletal muscle mass and function: Measurement and physiology of muscle fibre atrophy and muscle fibre loss in humans. Ageing Res Rev 2018;47:123–132.

23. Maden-Wilkinson TM, Degens H, Jones DA, McPhee JS. Comparison of MRI and DXA to measure muscle size and age-related atrophy in thigh muscles. J Musculoskelet Neuronal Interact 2013;13:320–328.

24. Andersen-Ranberg K, Petersen I, Frederiksen H, Mackenbach JP, Christensen K. Cross-national differences in grip strength among 50+ year-old Europeans: results from the SHARE study. Eur J Ageing 2009;6:227–236.

25. Ahrenfeldt LJ, Scheel-Hincke LL, Kjærgaard S, Möller S, Christensen K, Lindahl-Jacobsen R. Gender differences in cognitive function and grip strength: a cross-national comparison of four European regions. Eur J Public Health 2019;29:667–674.

26. Yeung SSY, Reijnierse EM, Trappenburg MC, Hogrel J-Y, McPhee JS, Piasecki M et al. Handgrip Strength Cannot Be Assumed a Proxy for Overall Muscle Strength. J Am Med Dir Assoc 2018;19:703–709.

27. Enoka RM, Christou EA, Hunter SK, Kornatz KW, Semmler JG, Taylor AM et al. Mechanisms that contribute to differences in motor performance between young and old adults. J Electromyogr Kinesiol 2003;13:1–12.

28. Carville SF, Perry MC, Rutherford OM, Smith ICH, Newham DJ. Steadiness of quadriceps contractions in young and older adults with and without a history of falling. Eur J Appl Physiol 2007;100:527–533.

29. Farina D, Negro F. Common synaptic input to motor neurons, motor unit synchronization, and force control. Exerc Sport Sci Rev 2015;43:23–33.

30. Castronovo AM, Mrachacz-Kersting N, Stevenson AJT, Holobar A, Enoka RM, Farina D. Decrease in force steadiness with aging is associated with increased power of the common but not independent input to motor neurons. J Neurophysiol 2018;120:1616–1624.

31. Challis JH. Aging, regularity and variability in maximum isometric moments. J Biomech 2006;39:1543–1546.

32. Hortobágyi T, DeVita P. Mechanisms Responsible for the Age-Associated Increase in Coactivation of Antagonist Muscles. Exerc Sport Sci Rev 2006;34.

33. Krishnan C, Allen EJ, Williams GN. Effect of knee position on quadriceps muscle force steadiness and activation strategies. Muscle Nerve 2011;43:563–573.

34. Smith SS, Woolley CS. Cellular and molecular effects of steroid hormones on CNS excitability. Cleve Clin J Med 2004;71 Suppl 2:S4–10.

35. Smith SS, Woodward DJ, Chapin JK. Sex steroids modulate motor-correlated increases in cerebellar discharge. Brain Res 1989;476:307–316.

36. Hall JE. Endocrinology of the Menopause. Endocrinol Metab Clin North Am 2015;44:485–496.

37. Dumitriu D, Rapp PR, McEwen BS, Morrison JH. Estrogen and the aging brain: an elixir for the weary cortical network. Ann N Y Acad Sci 2010;1204:104–112.

38. Waters EM, Mazid S, Dodos M, Puri R, Janssen WG, Morrison JH et al. Effects of estrogen and aging on synaptic morphology and distribution of phosphorylated Tyr1472 NR2B in the female rat hippocampus. Neurobiol Aging 2019;73:200–210.

39. Pandya M, Palpagama TH, Turner C, Waldvogel HJ, Faull RL, Kwakowsky A. Sex- and age-related changes in GABA signaling components in the human cortex. Biol Sex Differ 2019;10:5.

